# Simultaneous protein and RNA analysis in single extracellular vesicles, including viruses: SPIRFISH

**DOI:** 10.1101/2024.02.21.581401

**Authors:** Zach Troyer, Olesia Gololobova, Aakash Koppula, Zhaohao Liao, Felix Horns, Michael B Elowitz, Juan Pablo Tosar, Mona Batish, Kenneth W. Witwer

**Affiliations:** Department of Molecular and Comparative Pathobiology, Johns Hopkins University School of Medicine, Baltimore, Maryland, USA; EV Core Facility “EXCEL”, Institute for Basic Biomedical Sciences, Johns Hopkins University School of Medicine, Baltimore, Maryland, USA; Department of Medical and Molecular Sciences, and Department of Biological Sciences, University of Delaware, Newark, Delaware, USA; Howard Hughes Medical Institute and Division of Biology and Biological Engineering, California Institute of Technology, Pasadena, California, USA; Functional Genomics Laboratory, Institut Pasteur de Montevideo, Montevideo, Uruguay; School of Science, Universidad de la República, Montevideo, Uruguay; The Richman Family Precision Medicine Center of Excellence in Alzheimer’s Disease, Johns Hopkins University School of Medicine, Baltimore, Maryland, USA

**Keywords:** Single-particle, single-molecule, HIV, virus, extracellular vesicle, RNA, protein, smFISH, SP-IRIS

## Abstract

Interest in using nanoparticles for delivery of therapeutic RNA has been steadily growing, provoking a need to precisely understand their structure and contents. Single-particle and single-molecule analysis techniques provide snapshots of single biological nanoparticles, including viruses, liposomes, and extracellular vesicles (EVs). While existing methods primarily focus on protein detection, RNA delivery is becoming increasingly prevalent. A method to simultaneously detect protein and internal RNA in the same particle would reveal variability in size, structure, and RNA packaging efficiency, enabling optimization of nanoparticle delivery. Here, we introduce SPIRFISH, a high-throughput method for single-particle protein and RNA analysis, combining single particle interferometric reflectance imaging sensor (SP-IRIS) with single-molecule fluorescence in-situ hybridization (smFISH). Using SPIRFISH, we detect HIV-1 envelope protein and genomic RNA within single infectious virions, allowing resolution against EV background and noninfectious virions. We further show that SPIRFISH can be used to detect specific RNA within EVs. SPIRFISH should enable single particle analysis of a broad class of RNA-containing nanoparticles.

**Teaser:** A new single particle analysis technique simultaneously detects specific RNA and protein in biological nanoparticles.

## Introduction

Extracellular vesicles (EVs) have emerged as important mediators of intercellular communication and have garnered considerable attention in recent years (*1*, *2*). These nanoscale membrane-bound vesicles, released constitutively by various cell types, can encapsulate diverse cargo, including proteins, lipids, and nucleic acids (*3–5*). Their small size, propensity to interact with cells, and ability to protect internal cargos from the external environment due to the lipid bilayer has made EVs highly attractive as potential therapeutic agents (*6–8*). As such, there has been a considerable push to characterize the protein and nucleic acid content of EVs on a single particle basis (*9*, *10*).

EVs are produced by cells through various internal pathways and cellular machineries, giving rise to subsets of EVs that differ in size, contents, and function (*3–5*). Enveloped viruses are a subset of EVs in which a viral genetic program has hijacked the EV production machinery to release viral particles (virions), consisting of viral proteins and viral nucleic acids surrounded by a lipid-bilayer (*11–13*). Both virions and non-viral EVs are similar in size and host-protein content, as they share the same cellular origins, making their resolution as unique entities difficult in bulk analyses given that they are produced concurrently by the cell. Techniques to separate viruses from EVs by differing properties of density are often used, but are time consuming and may not always cleanly separate these populations (*13*). This difficulty is further compounded by the fact that virion production can be inefficient (*13*, *14*) producing a gradient of membranous particles that are part virus and part host yet lack the definitional replicative characteristic needed to be called a true infectious virus (e.g., an EV carrying a viral protein but not the viral nucleic acid). These challenges are bottlenecks in attempts to characterize surface proteins and internal cargos of true infectious virions.

In recent years, the field of single-particle analysis has witnessed significant advancements, enabling the multiparametric analysis of individual biological and synthetic particles with unprecedented precision (*9*, *10*, *15*). One such technique, known as single particle interferometric reflectance imaging sensor (SP-IRIS), has revolutionized the field by offering high-throughput analysis of nanoparticles captured by various antibody lawns on a silicon chip (*16–18*). SP-IRIS combines interferometric imaging with fluorescence microscopy, enabling size profiling, concentration, and protein biomarker analysis of individual biological particles within a bulk population. Addition of fluorophore-conjugated antibodies targeting surface or internal targets determines the presence or absence of up to 3 protein markers per particle; a 4^th^ marker is determined by the identity of the antibody capture lawn immobilizing the particle. SP-IRIS has mainly been applied to the study of EVs, but can theoretically be adapted to study of diverse biological nanoparticles, including viruses and liposomes.

A parallel technological advancement has allowed the development of single-molecule analysis techniques (*19*, *20*). Single-molecule fluorescence in situ hybridization (smFISH) has emerged as a powerful method for detecting and imaging individual RNA molecules within fixed cells and tissues (*21*, *22*). By using fluorescently labeled nucleic acid probes complementary to specific sequences, smFISH allows for the visualization and quantification of RNA at the single-molecule level. Typically, various 20-nucleotide probes are designed to bind adjacent regions of the target RNA molecule (*21*). The coincident binding of multiple probes results in a detectable fluorescent signal in a diffraction limited spot, which can be captured with conventional fluorescence microscopes. Since accumulation of multiple fluorophores on the target is needed to generate a detectable spot, sporadic, sequence-independent, off-target binding by a few probes generates a more diffuse signal. This provides very high signal to noise ratio and renders the technique highly specific (*21*).

An infectious enveloped virion cannot be defined by one characteristic alone. Coincident detection of multiple markers, both protein and RNA, is needed. Thus, we reasoned that the multiparametric single-particle analysis afforded by the SP-IRIS technique, in conjunction with fluorescent labeling of the viral genome by smFISH, would help identify a population of true infectious virions containing both viral envelope proteins for cellular entry and viral nucleic acid for replication. Furthermore, we reasoned that a combination of the smFISH and SP-IRIS techniques could be valuable for the EV field, in which the single-particle loading efficiency of therapeutic RNAs is often unknown (*23–25*). To our knowledge, no groups have successfully detected smFISH RNA signal within a single nanoparticle using SP-IRIS analysis.

In this study, we develop SPIRFISH as a novel approach that integrates the SP-IRIS technique with smFISH to enable the fluorescence-based detection of HIV-1 genomic RNA within single virions. First, we perform SP-IRIS analysis of replication competent HIV-1 virions, demonstrating that virions can potentially be distinguished from EVs by size and surface markers. We then perform SPIRFISH analysis to specifically detect fluorescent HIV-1 genomic RNA and identify infectious virions on a single-particle basis. We also examine extracellular vesicles generated by the COURIER system, which packages specific RNAs into extracellular vesicles using modified engineered protein nanocages (*26*). Finally, we demonstrate that the fluorescent RNA signal is reliant upon sequence complementarity of smFISH probes. This work serves as a proof-of-principle for the SPIRFISH workflow, and could be applied to specifically detect diverse nanoparticle-encapsulated RNAs and surface proteins in a single-particle, high-throughput manner.

## Results

### Production and characterization of HIV

Given the structural nature of the HIV lentivirus, consisting of an external protein-laden lipid bilayer surrounding a proteinaceous capsid containing two copies of viral single-stranded genomic RNA (gRNA) (*27*), we reasoned that HIV would serve as an ideal model system for which to develop the single-particle dual RNA and protein detection SPIRFISH methodology. Additionally, we were influenced by a particular issue within HIV research; HIV researchers have long recognized the need to analyze HIV-1 virions in the absence of “contaminating” EVs that are inevitably co-purified with viral preparations (*28*). The lack of available convenient methods for virion and EV separation has clouded research into the surface properties of true “infectious” virions – those carrying genomic RNA in addition to viral structural proteins (*29*). To analyze the single particle profile of HIV and develop the SPIRFISH methodology, we selected the CCR5-tropic HIV-1 strain BaL.

To produce highly concentrated, enriched HIV virions for analysis, chronically infected PM1-BaL cells were grown in cell culture and conditioned media collected during passaging. A large volume of virus-containing conditioned media was collected and then processed via a differential ultracentrifugation workflow (Figure 1A) designed to remove cells, cell debris, and large EVs. Conditioned media from uninfected PM1 cells was processed via the same workflow, in order to produce control PM1 EVs. As a negative control that accounts for any influence of EVs derived from the fetal bovine serum present in the RPMI formulation used with the PM1 cells, unconditioned media unexposed to cells was processed via the same workflow. The purified, concentrated virus from the 100,000g pellet was then characterized using a variety of methods. To validate that HIV virions were enriched via the collection workflow, a p24 ELISA detecting the HIV capsid structure was performed with the 100,000g pellet and the unprocessed conditioned media. The 100,000g pellet had ∼350X more concentrated p24 than a comparable volume of unprocessed conditioned media, confirming HIV virion enrichment (Figure 1B). To validate that the 100,000g pellet contains HIV virions, immunoblotting was performed against the HIV envelope protein gp120 and the HIV capsid protein p24 (Figure 1C). HIV Env gp120 was clearly visible as a single band, as expected. The p24 blot contained two bands, a more intense lower band corresponding to processed p24 and a fainter upper band corresponding to the unprocessed gag polyprotein p55 (typically stemming from immature viral particles). This pattern indicates that the majority of HIV virions purified in our workflow are in the mature, fully processed form.

**Fig. 1.**
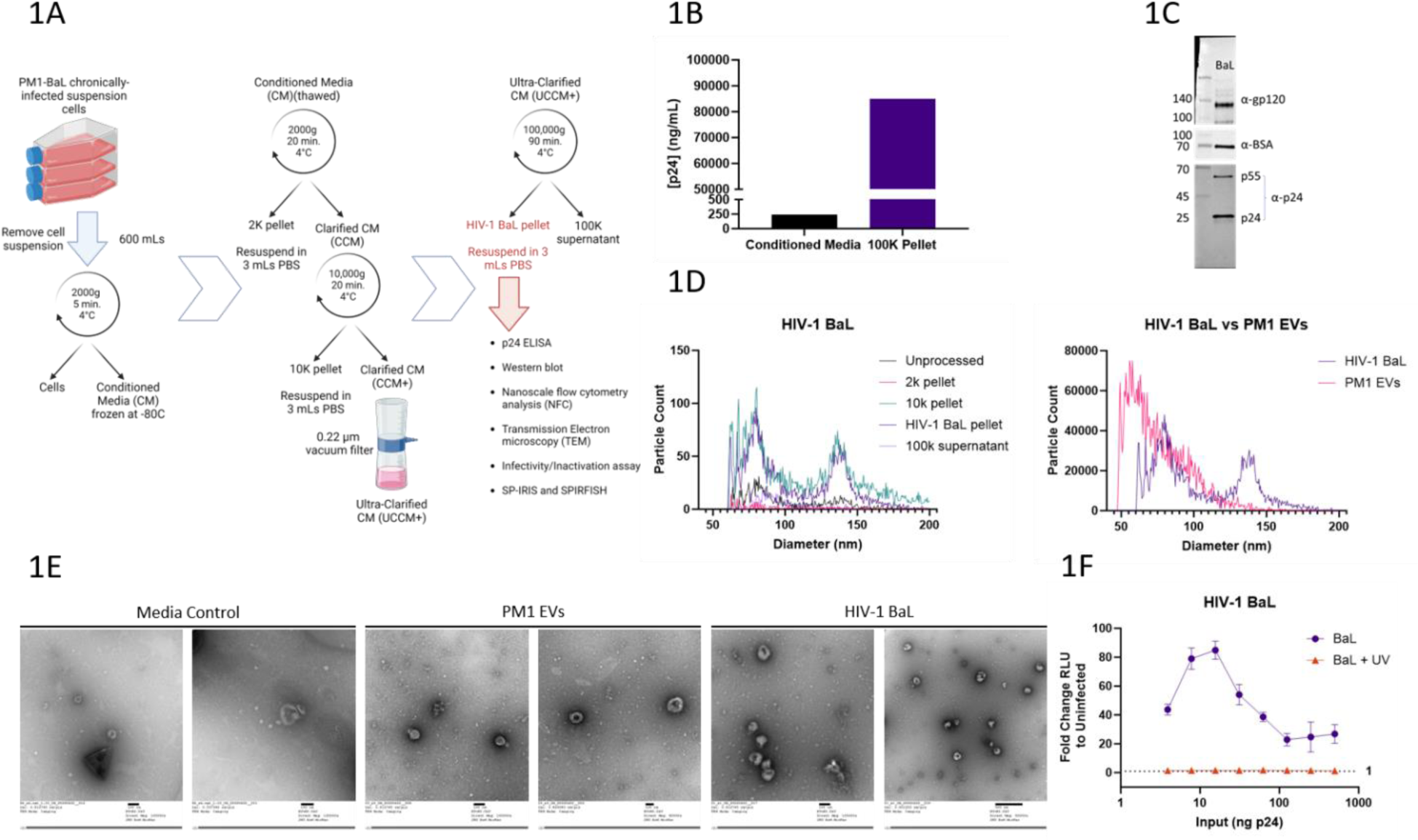
Production and characterization of HIV. **1A**: Schematic diagram depicting the differential ultracentrifugation workflow used to produce concentrated HIV for downstream analysis. **1B**: Bar graph showing the results of a p24 ELISA assay performed with unconcentrated HIV conditioned media and the concentrated HIV preparation. The differential ultracentrifugation workflow concentrated the HIV ∼350X. **1C**: Immunoblot analysis of the concentrated HIV preparation. HIV gp120 was visible as a single band ∼120 kDa. HIV Gag p24 was detected in two bands; a lower band (∼24 kDa) corresponding to the mature protein, and an upper band corresponding to unprocessed Gag polyprotein p55. Bovine serum albumin, a contaminant derived from FBS used in the cell culture media, was also detected. **1D**: **Left panel**: Nano-flow cytometry plots showing the size distribution of particles collected at various steps of the differential ultracentrifugation process. Different samples were run at different dilutions to avoid cytometer clogging, therefore particle counts represented on the Y-axis are not representative of true particle counts. **Right panel**: Nano-flow cytometry plots representing the concentrated HIV sample and the control PM1 extracellular vesicles (EVs). Samples were corrected for dilution factors before plotting. **1E**: Representative transmission electron micrographs of media control, PM1 EVs, and concentrated HIV. Individual scale bars are provided under each image. **1F**: Line chart representing the results of a TZM-Bl reporter luciferase assay measuring HIV infectivity at varying doses in the presence (orange line) or absence (purple line) of virus-inactivating UV-light. Results are represented as fold change compared to an uninfected cell control.

To characterize the size and concentration of the purified HIV, as well as further validate our viral enrichment, we ran the material collected from various stages of the collection workflow on a small particle flow cytometer capable of resolution down to 50 nm diameter (Figure 1D, left panel). Interestingly, the size histogram of the HIV enriched 100,000g pellet formed a distinct bimodal distribution, with one peak around 60-70 nm and a second around 140 nm in diameter. Based on the sizes of HIV virions reported in the literature, it is highly likely that the second, larger size peak represents intact HIV virions, although the contribution of non-viral EVs to this peak cannot be excluded (*30*, *31*). The smaller peak likely consists mostly of EVs, as the viral capsid structure creates a lower size limit for intact virions. Supporting these conclusions, a size histogram of the PM1 EVs from uninfected cells was conspicuously missing the larger size peak, while strongly overlapping with the lower size peak seen in the HIV sample (Figure 1D, right panel). The nanoflow data also supported the enrichment of the HIV virions via our workflow; the 100,000g pellet contained the highest particle concentration and the most clearly visible larger size peak. Of note, there is some evidence of HIV virions in the 10,000g pellet, as a bimodal size distribution was also seen. These could represent viral aggregates (*32*). Transmission electron microscope (TEM) images of PM1 EV and HIV 100,000g pellets revealed typical cup-shaped, collapsed EV morphologies, an artifact of the negative staining TEM preparation (Figure 1E) (*33*). The HIV TEM images contained particles with visibly electron dense interiors, potentially indicating presence of the HIV virions.

To test the infectivity of the purified HIV, we utilized the TZM-Bl reporter assay (*34*). TZM-Bl cells are a HeLa cell derivative that produce firefly luciferase under control of the HIV trans-activator of transcription (Tat) protein, which is produced during productive viral infection. The purified HIV was infectious over a broad range of doses, although the highest levels of productive infection were found at a medium dose, likely due to cellular toxicity from higher doses interfering with luciferase production (Figure 1F). Importantly, purified HIV could be rendered non-infectious at all tested doses via a 20-minute UV light exposure. Inactivation is necessary for safe handling of virus for SP-IRIS and SPIRFISH in the following sections. Although UV-inactivation is thought to work primarily by damaging the viral RNA genome, it does so through RNA-protein crosslinking and formation of pyrimidine dimers which should not affect the genomic sequence (*35*).

### SP-IRIS Analysis of HIV reveals identifiable virions that can be distinguished from EVs

After standard bulk characterization of our HIV preparation, we proceeded to performing multi-parametric single particle characterization of HIV virions. For this type of analysis, we used single-particle interferometric reflectance imaging sensor (SP-IRIS) technology, as it can report size and multiple protein surface markers for each particle detected (*16–18*). We hypothesized that HIV virions would be captured by the tetraspanin capture antibodies on the SP-IRIS chips, due to the presence of tetraspanins in the HIV lipid envelope (*36*). Additionally, we hypothesized that HIV virions on the SP-IRIS chips would be distinguishable from the EVs within the same sample due to increased particle diameter and presence of the HIV-1 Env protein gp120 (detected by fluorescent antibody during the SP-IRIS scanning phase). UV-inactivated media control, PM1 EVs, and HIV were incubated overnight with SP-IRIS chips, probed with fluorescent antibodies, and then scanned. The total particle capture across the three anti-tetraspanin antibody capture spots on the SP-IRIS chips was compared across conditions (Figure 2A, left panel). As expected, the media control had a small amount of particle capture, likely consisting of FBS-derived EVs (*37*). The PM1 EVs and the HIV sample had higher levels of particle capture than the media control, and were relatively similar. The HIV sample had slightly more particles captured by the CD63 spot as compared to the PM1 EVs. The particles captured on each tetraspanin spot were further broken down into subcategories based on the incidence of fluorescence detection by antibodies targeting surface proteins CD63, CD81, and HIV-1 Env gp120 on a per-particle basis (Figure 2A, right panels). Fluorescent events detected in the media control were primarily gp120+, and occurred primarily in the CD9 and CD63 capture spots – likely antibody background. Interestingly, both PM1 EVs and HIV had high levels of particle capture on the CD81 spot. For the PM1 EVs, most of these particles were CD81+_capture_/CD81+_fluorescence_, with a smaller fraction of CD81+_capture_/CD63+_fluorescence_ particles and almost no CD81+_capture_/gp120+_fluorescence_ particles. For HIV, there was a markedly different distribution. In addition to a large amount of CD81+_capture_/CD81+_fluorescence_ particles, there were also a large amount of CD81+_capture_/CD63+_fluorescence_ particles. Importantly, there was a novel population of CD81+_capture_/gp120+_fluorescence_ particles not seen in the media control or PM1 EVs – likely gp120+ HIV virions that have incorporated the tetraspanin CD81.

**Fig. 2.**
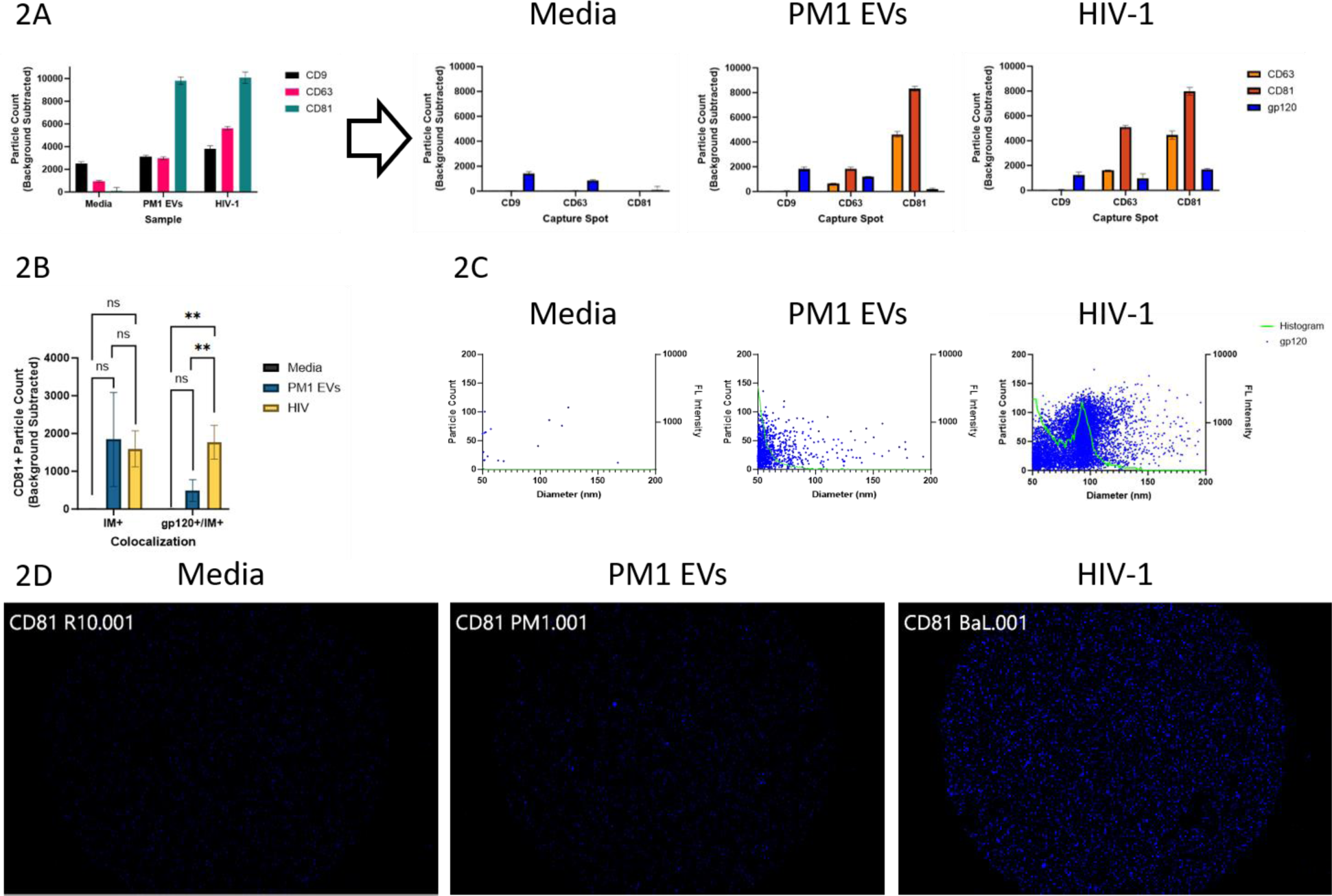
SP-IRIS Analysis of HIV Virions. **2A**: **Left panel**: Bar graph comparing total particle capture of media control, PM1 extracellular vesicles (EVs), and HIV across different anti-tetraspanin capture spots on SP-IRIS microchips. Graph is from one biological replicate and is representative of the trend seen across experiments. Bars show mean particle counts across three replicate capture spots on the same SP-IRIS microchip; error bars represent standard deviation. **Right panels**: Bar graphs indicating the number of particles on each antibody capture spot measured as positive for a target protein. Bars show mean particle counts across three replicate capture spots within an SP-IRIS microchip; error bars represent standard deviation. **2B**: Grouped bar graph comparing number of CD81+ particles detected by interferometry, or detected by interferometry and gp120+. Bars show mean particle counts across three independent SP-IRIS experiments; error bars represent standard deviation. Statistical testing was performed by one-way ANOVA with Tukey’s multiple comparisons test, within each category (for representation, data are within a single graph). ns, P > 0.05; *, P ≤ 0.05; **, P ≤ 0.01; ***, P ≤ 0.001. **2C**: Combined representative size histogram/fluorescent scatter plots of particles captured on the CD81 capture spots. Green line represents size distribution of particles detected interferometrically, with size plotted on X-axis and particle count on left Y-axis. Blue dots represent gp120+ particles co-detected interferometrically, plotted with size on X-axis and gp120 fluorescence intensity on right Y-axis. **2D**: Representative blue channel (gp120+) fluorescent images of CD81 capture spots on SP-IRIS microchips.

Since HIV virions were being captured by CD81, we decided to focus on this antibody capture spot. Given that our previous size analysis had indicated that our HIV sample has a population of larger particles, likely virions, we decided to add a size parameter to our analysis. To do this, we sorted the data to only view particles that had been detected and sized interferometrically by SP-IRIS (fluorescence events can be detected without an associated size measurement). PM1 EVs and HIV samples both had higher levels of IM+/CD81+_capture_/gp120-_fluorescence_ particles as compared to the media control, although the differences were not statistically significant (p=0.0599 and p=0.0983, respectively) due to variability in capture on chips run on different days (Figure 2B, IM+). This population is likely to be non-viral EVs. The HIV sample had a significantly higher level of IM+/CD81+_capture_/gp120+_fluorescence_ particles – likely virions – as compared to the PM1 EVs (p=0.0054) and media control (p=0.0010). Within this colocalization category, the PM1 EVs were statistically indistinguishable from the media control (p=0.2104).

To further investigate if virions are distinguishable due to larger size within the IM+/CD81+_capture_ particles in the HIV sample, we plotted the interferometric size distribution as a size histogram (Figure 2C). Like our small particle flow cytometer analysis, a distinct bimodal distribution was observed for the HIV sample that was not present for the PM1 EVs. Interestingly, the HIV sample’s larger SP-IRIS size peak (IM_High_) centered over ∼95 nm, differing from the ∼140 nm seen with the small particle flow cytometer. This may be due to the different methods used by these techniques to ascertain particle size. To determine if particles that made up the larger size peak are gp120+ as would be expected of infectious HIV virions, we plotted each IM+/CD81+_capture_/gp120+_fluorescence_ particle on the size histogram as a function of IM size (X-axis) and gp120 fluorescence intensity (right Y-Axis). The majority of gp120+ particles in the HIV sample were also IM_High_. Some gp120+ particles were seen in the smaller size peak for HIV; these were also seen in the PM1 EV’s size peak and thus likely represent EVs nonspecifically binding gp120 antibody. Importantly, the IM_High_ gp120+ particles corresponding to the second size peak in the HIV sample also had higher gp120 fluorescence intensities (gp120_high_) as compared to the smaller gp120+ particles seen in the HIV and PM1 EV samples. This brighter fluorescence was also seen visually when viewing fluorescent images of the circular CD81 capture spots on the various samples’ SP-IRIS microchips (Figure 2D). Taken together, these results suggest that potential HIV virions can be identified and distinguished from EV background as CD81+_capture_/IM_High_/gp120_high_ particles.

### Design and validation of HIV genome-targeting single-molecule fluorescence in-situ hybridization (smFISH) probes

Our SP-IRIS results so far have revealed a novel population of CD81+_capture_/IM_High_/gp120_high_ particles present only in the HIV sample. While it would be tempting to say that these are authentic HIV virions, such an assertation is not yet supported by these data. It is well known that non-infectious EVs can carry viral envelope glycoproteins, Including that of HIV (*38*, *39*). It remains possible that some CD81+_capture_/IM_High_/gp120_high_ particles we detect in the HIV sample were actually large EVs carrying gp120+ but having no viral gRNA and no replicative capacity. Infectious virions are therefore expected to contain both gp120 and HIV gRNA. In order to fluorescently detect HIV gRNA, we decided to use single-molecule fluorescence in-situ hybridization (smFISH) (*21*, *22*). In this technique, multiple fluorescently-labeled short DNA probes collectively bind a complementary RNA sequence yielding a diffraction limited spot that represent individual RNA molecules. A pool of 219 smFISH probes were designed to target complementary regions spanning the ssRNA genome of the HIV strain BaL (Figure 3A, 3C, Data S1). To confirm that these smFISH probes bind specifically to HIV BaL gRNA, uninfected PM1 and chronically infected PM1-BaL cells, used to make the control PM1 EVs and HIV preparations respectively, were grown on glass coverslips, fixed, and hybridized using an smFISH hybridization workflow and Texas Red-labeled probes. Only PM1-BaL cells were visibly fluorescent with Texas Red, as expected since active viral replication was occurring in these cells (Figure 3B). Uninfected PM1 cells did not have any Texas Red fluorescence, demonstrating the specificity of the smFISH probe set for the HIV-1 BaL gRNA.

**Fig. 3.**
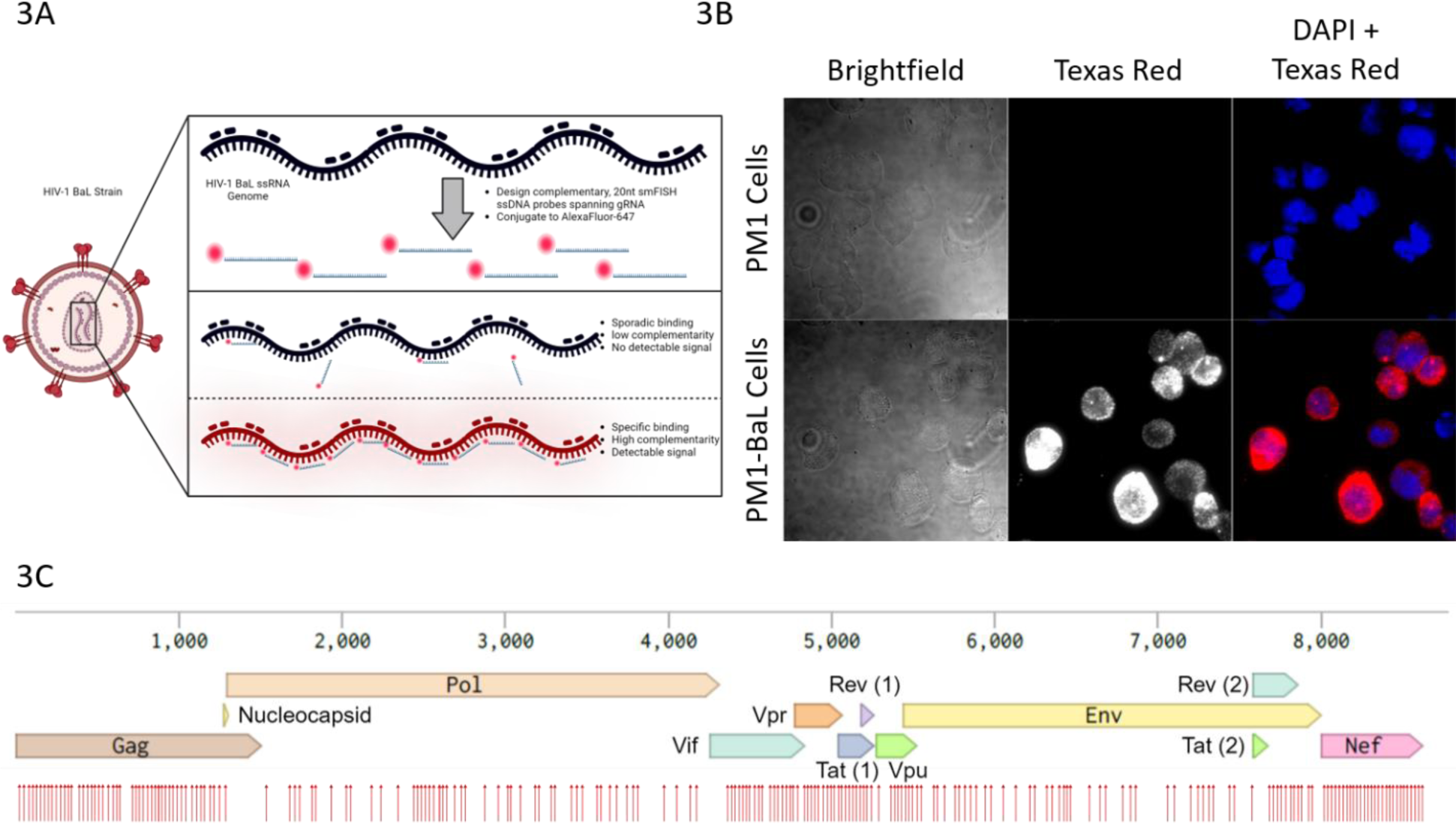
Design and Validation of HIV-1 BaL smFISH Probes. **3A**: Schematic representation of the principal of smFISH and how smFISH probes were designed complementary to the HIV-1 BaL genome. **3B**: Fluorescent microscopy images of slides containing cells hybridized with the HIV-1 BaL probes via conventional smFISH. 100X Magnification. The Texas Red-labelled probes do not hybridize with uninfected PM1 cells (top middle panel). HIV-1 BaL infected PM1 cells are hybridized with smFISH probes due to the presence of HIV RNA in the cytosol, resulting in Texas Red signal (bottom middle panel – represented in greyscale). Nuclei are visible due to DAPI staining (right panel – top and bottom). **3C:** Schematic of the HIV-1 BaL genome, with genes indicated by text and block arrows. The binding locations of individual probes within the HIV-1 BaL gRNA smFISH probe pool are represented by the red arrows.

### SPIRFISH Analysis of HIV identifies and distinguishes infectious virions from noninfectious virions and EVs

Building upon our SP-IRIS analysis of HIV, we reasoned that we could combine smFISH fluorescent-probe hybridization with the fluorescent-antibody protein detection step of the SP-IRIS workflow. This would theoretically allow coincident fluorescence-based detection of specific protein and internal RNA on a single particle basis; a modified SP-IRIS/smFISH workflow we call SPIRFISH. In this case we attempted to use SPIRFISH for the simultaneous detection of HIV gp120 and gRNA, to further distinguish infectious viral particles from particles or defective virions lacking one or both of these viral elements. Similar to our previous SP-IRIS experiments, UV-inactivated media control, PM1 EVs, and HIV were incubated with SP-IRIS chips overnight, probed/hybridized with fluorescent antibodies and HIV gRNA smFISH probes, and then scanned. The total particle capture across the three anti-tetraspanin antibody capture spots on the SP-IRIS chips was compared across conditions (Figure 4A, left panel). The results were similar to those from our previous SP-IRIS analysis; the media control had very small amount of capture, while PM1 EVs and HIV had similar higher levels of capture with most particles captured by the CD81 spot. The captured particles on each antibody spot were again broken down into subcategories based on fluorescence detection of HIV gp120, CD81, and hybridized smFISH probes targeting HIV gRNA (Figure 4A, right panels). In terms of gp120 and CD81 fluorescence, results were largely similar to our previous SP-IRIS analysis, particularly within the CD81 capture spot. Importantly, there was a new population of CD81+_capture_/smFISH+_fluorescence_ particles in the HIV sample that was not seen in the control samples. It is possible that these are infectious HIV virions in which the viral gRNA was being fluorescently detected during SPIRFISH scanning (Figure 4A, right panel, HIV-1). Surprisingly, this new population of smFISH+ particles was detected at similar levels whether or not a permeabilization-based washing protocol was used for the chip (data not shown).

**Fig. 4.**
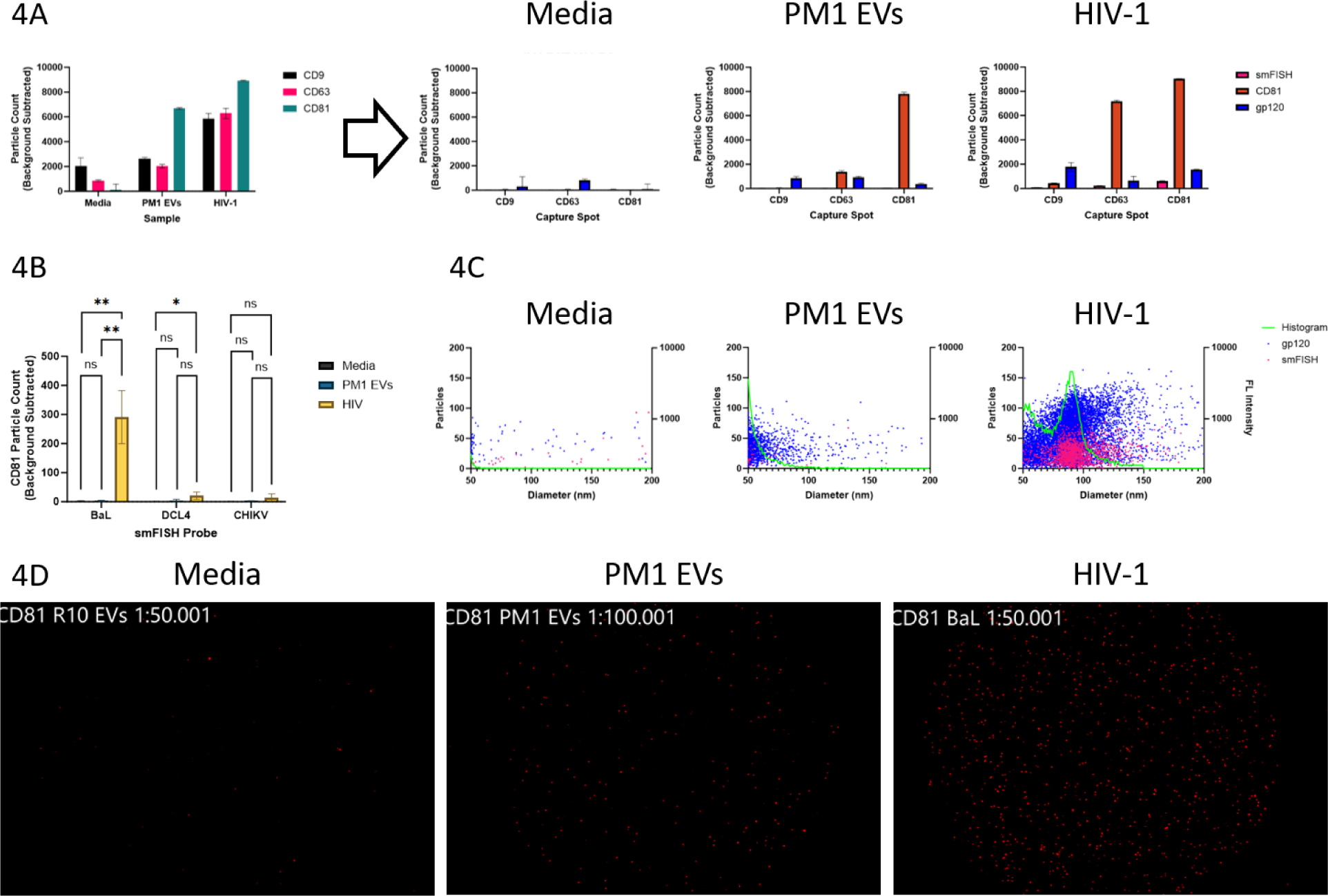
SPIRFISH Analysis of HIV Virions. **4A**: **Left panel**: Bar graph comparing particle capture of media control, PM1 extracellular vesicles (EVs), and HIV virions across different anti-tetraspanin capture spots on SP-IRIS microchips. Graph is from one biological replicate and is representative of trend seen across experiments. Bars show mean particle counts across three replicate capture spots on the same SP-IRIS microchip; error bars represent standard deviation. **Right panels**: Bar graphs indicate the number of particles on each antibody capture spot measured as positive for a target protein or HIV gRNA. Bars show mean particle counts across three replicate capture spots on the same SP-IRIS microchip; error bars represent standard deviation. **4B**: Grouped bar graph showing number of CD81+/gp120+/smFISH+ particles co-detected interferometrically. Bars show mean particle counts across three independent SP-IRIS experiments; error bars represent standard deviation. Statistical testing was performed by one-way ANOVA with Tukey’s multiple comparisons test, within each category (for representation, data are within a single graph). ns, P > 0.05; *, P ≤ 0.05; **, P ≤ 0.01. **4C**: Combined representative size histogram/fluorescent scatter plots of particles captured on CD81 capture spots. Green line represents size distribution of particles detected interferometrically, with size plotted on X-axis and particle count on left Y-axis. Blue (gp120+) or magenta (smFISH+) dots represent fluorescent particles co-detected interferometrically, plotted with size on X-axis. Fluorescence intensities for gp120 and smFISH signals are represented on right Y-axis. **4D**: Representative red channel (HIV-1 gRNA smFISH+) fluorescent images of CD81 capture spots on SP-IRIS microchips.

As with our previous SP-IRIS analysis, we added a particle size component to the SPIRFISH data given our previous observation that likely HIV virions tend to be larger and therefore IM+. To look closer at the novel smFISH+ population that we suspect are infectious HIV virions, we analyzed the number of IM+/CD81+_capture_/gp120+_fluorescence_/smFISH+_fluorescence_ particles across the different conditions (Figure 4B). There were close to zero particles matching these parameters in the media control and PM1 EV conditions; comparatively, there were a significant amount of these particles in the HIV condition (p=0.0013 vs both PM1 EVs and media control). These results lend further support to the idea that IM+/CD81+_capture_/gp120+_fluorescence_/smFISH+_fluorescence_ particles are infectious virions, and suggest that smFISH probes are hybridizing with and labeling HIV gRNA. Interestingly, the IM+/CD81+_capture_/gp120+_fluorescence_/smFISH+_fluorescence_ particles represent 13.78% of the total IM+/CD81+_capture_/gp120+_fluorescence_ population, suggesting approximately 1 in 8 captured virions carry genomic RNA (Data S2). To rule out the possibility that the smFISH probes are non-specifically binding to HIV virions, we used two non-HIV specific smFISH probe sets – one targeting DCL4 mRNA from *Arabidopsis thaliana* and another targeting the Chikungunya virus (CHIKV) nsP2 gene. When SPIRFISH was performed using the CHIKV smFISH probe set, the population of IM+/CD81+_capture_/gp120+_fluorescence_/smFISH+_fluorescence_ in the HIV sample was reduced to near background levels indistinguishable from non-viral controls. With the DCL4 probe set, this same population was also reduced to near background levels in the HIV-1 sample, although a small amount of non-specific binding was present (p=0.0374) lending a statistical increase as compared to media control. These results suggest that smFISH probes are capable of being used in SPIRFISH analysis to specifically detect target RNA in conjunction with protein in single particles.

In our SP-IRIS analysis of HIV, we concluded that HIV virions are likely CD81+_capture_/IM_High_/gp120_high_ due to propensity for gp120+ particles, especially those with high fluorescent intensity, to fall within the IM_High_ secondary size peak when plotted on a size histogram. To determine if smFISH+ particles preferentially fall within the IM_High_ size peak, as would be expected of infectious virions carrying HIV gRNA, we plotted each IM+/CD81+_capture_/smFISH+_fluorescence_ particle on the size histogram as a function of IM size (X-axis) and smFISH fluorescence intensity (right Y-Axis) (Figure 4C). We also plotted the IM+/CD81+_capture_/gp120+_fluorescence_ particles in the same way, to see if the population overlaps. Indeed, the smFISH+ particles almost universally fall into the IM_High_ size peak, alongside the majority of the gp120_high_ particles. Very few smFISH+ particles are found in the IM_Low_ size peak that we believe contains primarily EVs. Interestingly, the smFISH fluorescence intensity in the HIV sample was relatively low compared to gp120 fluorescence, perhaps reflecting differing molecular target availability. Nevertheless, the smFISH signal was bright enough to be clearly distinguishable from the control samples when viewing fluorescent images of the circular CD81 capture spots on the SP-IRIS microchips (Figure 4D). Taken together, these data suggest that infectious HIV virions can be distinguished from noninfectious virions and EVs using SPIRFISH analysis as CD81+_capture_/IM_High_/gp120_high_/HIV gRNA smFISH+ particles.

### SPIRFISH detection of EV-encapsulated tRNA

We aimed to validate the applicability of our SPIRFISH technique to extracellular vesicles, with a specific focus on the detection of tRNAs and tRNA fragments due to their high abundance in EVs (*40–42*). In particular, our investigation centered on the detection of 5’ tRNA^Gly^GCC, which exhibited elevated abundance in EVs, as reported (*41*). Specifically, we expected to detect nicked forms and fragments derived from 5’ tRNA^Gly^GCC, due to their stability (*40*) and increased abundance in EVs (*42*).

To achieve this, we designed a set of fluorescently-labeled short DNA probes, akin to those employed in HIV gRNA smFISH. These probes were tailored to target complementary regions within 5’ tRNA^Gly^GCC sequences commonly found in EVs, covering full-length tRNA, nicked forms, or tRNA fragments. Our experimental approach involved incubating purified EV samples on SP-IRIS chips, followed by fixation, permeabilization, and hybridization with tRNA-specific smFISH probes and fluorescent antibodies. This protocol, similar to that used for HIV gRNA detection but with the additional fixation and permeabilization steps, was applied to EVs from two different cell culture lines: U-2 OS and Expi293F.

Figure 5A depicts the total particle distribution captured on the tetraspanins printed on the chip. In Figure 5B, the distribution of fluorescent signals for smFISH probe (5’ tRNA^Gly^GCC), CD81, and CD9 across different capture spots is presented for both cell lines. As anticipated, the tRNA signal was low compared to the protein signals but remained slightly above background levels. Examination of colocalization between tetraspanins and the smFISH probe (Figure 5C) revealed that the majority of tRNA was present in particles that were CD9+ and CD81+ (triple positive EVs) for U-2 OS, while in expi293F, tRNA colocalized with CD81+ EVs captured on CD9 or CD81 spots. DCL4 smFISH probe, used as a negative control, showed lower signal compared to tRNA signal for both cell lines.

**Fig. 5.**
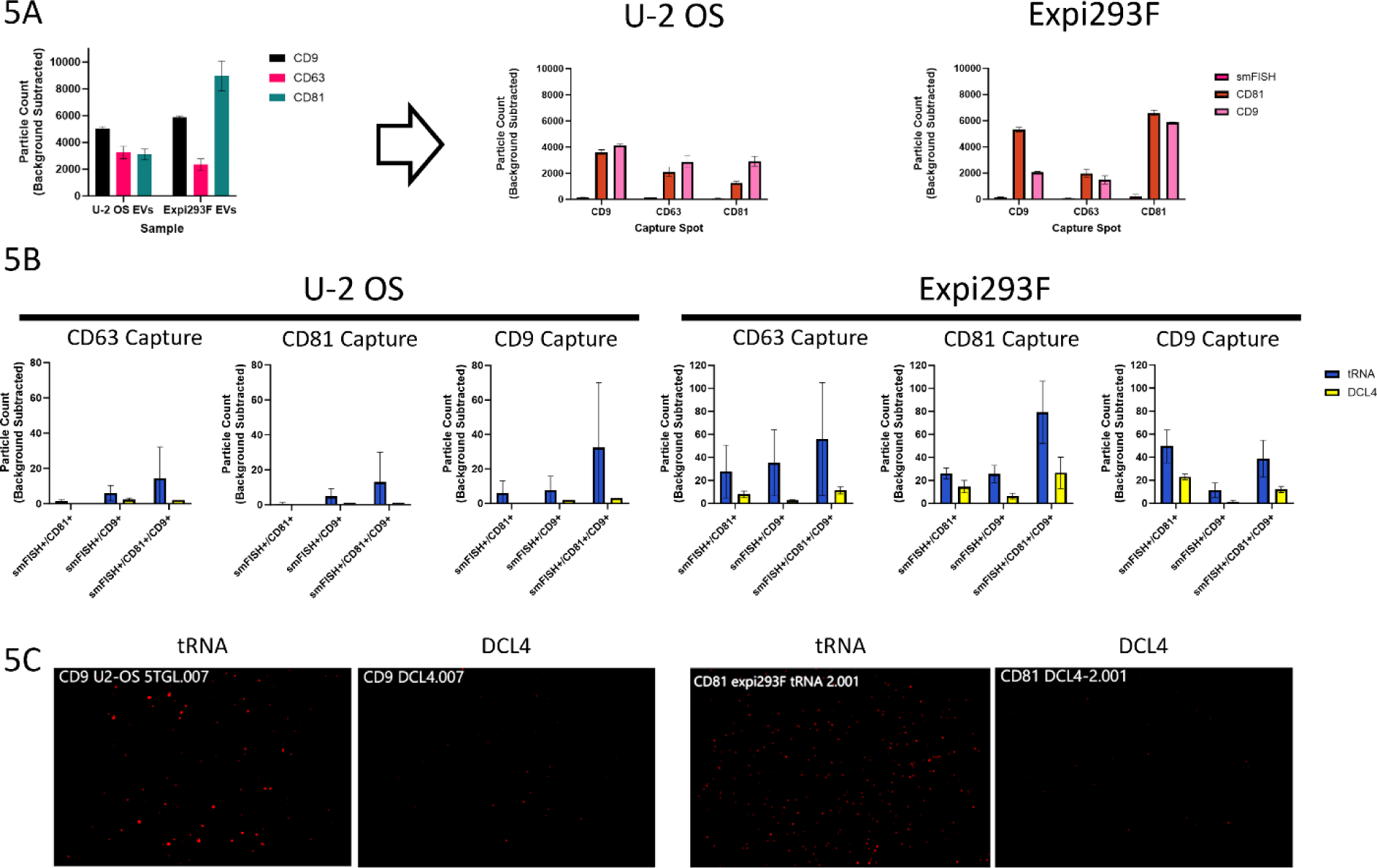
SPIRFISH detection of EV-encapsulated tRNA. **5A**: **Left panel**: Bar graph comparing total particle capture of U-2 OS and Expi293F extracellular vesicles (EVs) across three anti-tetraspanin capture spots on SP-IRIS microchips. Graph is from one biological replicate and is representative of trend seen across experiments. Bars show mean particle counts across three replicate capture spots on the same SP-IRIS microchip; error bars represent standard deviation. **Right panels**: Bar graphs indicating number of particles on each anti-tetraspanin antibody capture spot measured as positive for a target protein or 5’ tRNA^Gly^GCC. Bars show mean particle counts across three replicate capture spots (or four for Expi293F/tRNA condition) on the same SP-IRIS microchip; error bars represent standard deviation. **5B**: Various grouped bar graphs showing breakdown of smFISH+ particle counts by single-particle protein marker colocalization on each anti-tetraspanin antibody capture spot, separated by use of specific (blue bars) or non-specific (yellow bars) smFISH probes. Data from U-2 OS EVs is shown on the left and from Expi293F EVs on the right. Bars show the mean particle counts across three independent SP-IRIS microchip experiments; error bars represent standard deviation. **5C**: Representative red channel (5’ tRNA^Gly^GCC smFISH+) fluorescent images of the CD9 capture spot (U2-OS) and CD81 capture spot (Expi293F) on the SP-IRIS microchips, for both EV types and from the specific and non-specific smFISH probe experiments. Brightness was adjusted by +40% for all images to increase visibility of smFISH+ fluorescence.

Figure 5D displays raw fluorescence images of the spots, indicating that the 5’ tRNA^Gly^GCC signal is brighter and more abundant than that of DCL4 for both cell lines. The tRNA smFISH signal, being less abundant and dimmer than that measured for HIV virions, requires optimization of background correction to reduce nonspecific signals. In our approach, we subtracted the individual control IgG spot capture from the tetraspanin capture to achieve this correction.

### SPIRFISH detection of mCherry mRNA in engineered EVs

As a final demonstration of the versatility of SPIRFISH, we obtained engineered EVs designed to incorporate mCherry mRNA via a novel enveloped protein nanocage (EPN) system, EPN24-MCP (*26*). These EVs contain self-assembling 60-subunit nanocages possessing inward facing MS2 bacteriophage coat protein (MCP) motifs, which bind to cognate MS2-loops engineered in mRNA, (*43*) facilitating mRNA loading of the nanocage. Incorporation of the HIV p6 peptide secretion signals allows self-assembled, RNA-containing nanocages to be released inside EVs. EPN structures can also incorporate EGFP protein due to readthrough of a T2A-EGFP ribosome skip site downstream of the EPN structural component within the EPN plasmid. We obtained two variants of EPN24-MCP EVs: one designed to package Cre Recombinase mRNA and another for mCherry mRNA.

We performed SPIRFISH using the two types of EPN24-MCP EVs and an smFISH probe set targeting mCherry mRNA. As we expected, the SP-IRIS microchips used in the SPIRFISH analysis were able to capture abundant EPN24-MCP EVs on all three of the anti-tetraspanin capture spots (Figure 6A, left panel). Both Cre mRNA and mCherry mRNA EPN24-MCP EVs had similar patterns of capture across the three tetraspanin spots: fewer particles captured by CD9 and more captured by CD63 and CD81. The mCherry mRNA EVs had slightly higher particle capture than the Cre mRNA EVs, overall. The particles captured on each tetraspanin spot were further broken down into subcategories based on fluorescence detection of mCherry mRNA and EPN EGFP (Figure 6A, right panels). The EGFP contained within the EPN structure was readily detectable on all three capture spots for both EV subtypes. As expected, the signal originating from the smFISH probes targeting mCherry mRNA was only detectable in the mCherry mRNA EPN24-MCP EVs, demonstrating probe specificity. This smFISH signal was readily visible as bright red fluorescence on the images of the SP-IRIS microchip capture spots (Figure 6D). Only low background levels of smFISH signal were detected in the Cre mRNA EPN24-MCP EVs. For the mCherry mRNA EPN24-MCP EVs, mCherry mRNA smFISH signal was seen in CD9+, CD63+, and CD81+ EVs – in proportions mirroring the total capture and the EGFP signal.

**Fig. 6.**
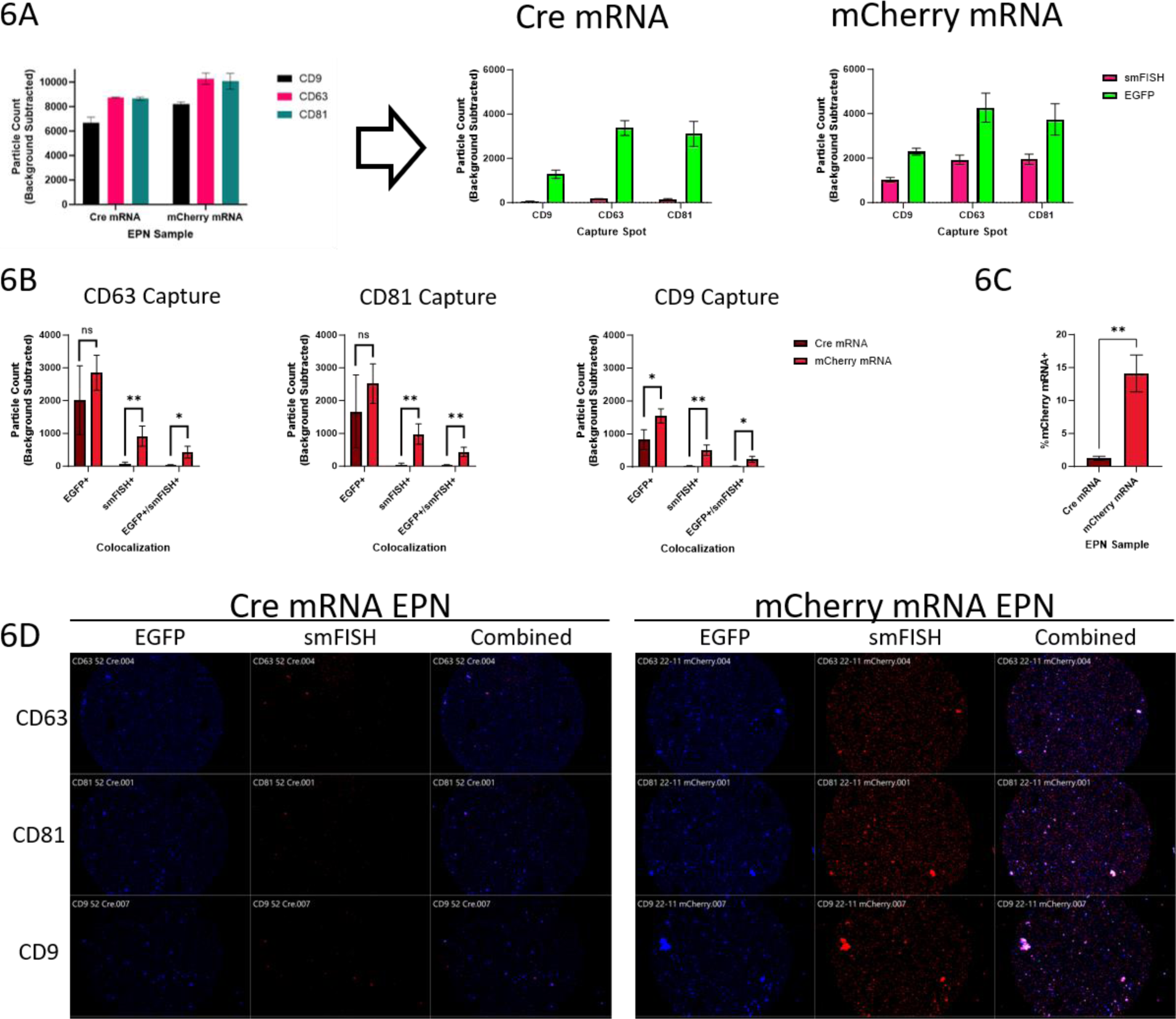
SPIRFISH Analysis of Engineered EVs. **6A**: **Left panel**: Bar graph comparing particle capture of extracellular vesicles (EVs) designed to package Cre or mCherry mRNA, across three anti-tetraspanin capture spots on SP-IRIS microchips. Graph is from one biological replicate and is representative of trend seen across experiments. Bars show mean particle counts across three replicate capture spots on the same SP-IRIS microchip; error bars represent standard deviation. **Right panels**: Bar graphs indicating number of particles on each capture spot measured as EGFP+ or mCherry mRNA+. Bars show mean particle counts across three replicate capture spots on the same SP-IRIS microchip; error bars represent standard deviation. **6B**: Grouped bar graphs showing particle counts by EGFP/mCherry mRNA colocalization on each capture spot, separated by Cre-mRNA (brown bars) or mCherry-mRNA (red bars) EVs. Bars show mean particle counts across three independent SP-IRIS experiments; error bars represent standard deviation. Statistical testing comparing counts between Cre-mRNA EVs and mCherry-mRNA EVs was performed using unpaired two-tailed T-test, within each category (for representation, data are within a single graph). ns, P > 0.05; *, P ≤ 0.05; **, P ≤ 0.01. **6C**: Bar chart indicating percentage of smFISH+ particles on SP-IRIS microchips. Bars show mean particle counts across three independent SP-IRIS microchip experiments; error bars represent standard deviation. Statistical testing was performed using unpaired two-tailed T-test. ns, P > 0.05; *, P ≤ 0.05; **, P ≤ 0.01. **6D**: Fluorescent images of different capture spots on SP-IRIS microchips. Left column (EGFP), middle (mCherry mRNA), right (combined).

We then performed colocalization analysis within each capture spot to determine how many EVs were double positive for EPN-EGFP and the mCherry mRNA smFISH signal (Figure 6B). The number of EGFP+ single positive EVs was similar on the CD63 and CD81 capture spots between the two EV conditions, suggesting equal amounts of EPN24-MCP incorporation. A small difference in EGFP+ single positive EVs was seen on the CD9 capture spot (p=0.0282). There were extremely few smFISH+ single positive EVs within the Cre mRNA condition, irrespective of capture spot, as these EVs do not possess requisite mCherry mRNA to generate smFISH signal above background levels. Comparatively, there were a significantly higher (∼10-20 fold) number of smFISH+ single positive EVs within the mCherry mRNA condition on all three capture spots (CD63: p=0.0094; CD81: p=0.0070; CD9: p=0.0074). These events could be EVs lacking the EPN structure but incorporating mCherry mRNA randomly, or EVs containing mCherry mRNA and EPN24-MCP for which the T2A facilitated ribosome skipping and avoided EGFP structural incorporation. The number of double positive EPN-EGFP+/mCherry mRNA+ EVs followed a similar pattern, with significantly higher numbers in the mCherry mRNA condition for all three capture spots (CD63: p=0.0195; CD81: p=0.0086; CD9: p=0.0189). Finally, we determined the EV loading efficiency of mCherry mRNA, a useful metric for therapeutic particle development, by calculating the percentage of total captured particles that were smFISH+. The mCherry mRNA EPN24-MCP EVs had an average loading efficiency of 14.1148%, significantly higher than background levels (p=0.0014); about 1 in 7 engineered EVs carry mCherry mRNA by this metric (Figure 6C).

## Discussion

The simplicity, sensitivity, and the ability to multiplex single-particle protein and RNA detection afforded by SPIRFISH makes it a powerful analysis technique that can be used to unravel population-level heterogeneity of diverse nanoparticles. SPIRFISH can be used with many biological nanoparticles that are currently under consideration for their use as therapeutics vectors; this includes extracellular vesicles (EVs), liposomes, and virus-like particle (VLPs) or intact viruses (*6*, *44–47*). Often, these particles are engineered and manipulated to package various proteins and RNA molecules with therapeutic functions. Given new appreciation of nanoparticle-population heterogeneity and burgeoning single particle analysis techniques (*9*, *10*, *15*, *48*), SPIRFISH fills a niche for single-particle biomarker colocalization analysis where bulk analysis methods do not suffice (*49–52*). Furthermore, its simultaneous detection of RNA and protein are an improvement over current colocalization analysis methods, which mostly focus on proteins due to the simplicity of antibody binding (*16–18*, *53*). Given the increasing focus on packaging RNA species into nanoparticles, including mRNAs, miRNAs, and siRNAs, there is a clear need to detect RNA on a single-particle basis (*6*, *54*, *55*). Moreover, in some cases colocalization of the therapeutic RNA with a surface protein intended to improve RNA delivery is important for particle functionality. A measurement of the percent of single particle RNA/protein colocalization in such a situation could reflect the therapeutic efficacy of the bulk nanoparticle population.

Reflecting the aforementioned need for single-particle RNA detection, a recent study developed a new technique called single-EV and particle protein and RNA assay (^siEVP^PRA) and successfully detected colocalized single-particle protein and RNA (*56*). This technique immobilizes particles on micropatterned, functionalized cover glass surfaces using biotinylated capture antibodies, before probing with fluorescent antibodies and customized fluorescent molecular beacons targeting RNA or mRNA subregions. Antibodies and molecular beacons were added sequentially and at different temperatures, before imaging with TIRF microscopy at high magnification. Similar to ^siEVP^PRA, SPIRFISH is also capable of detecting colocalized single-particle protein and RNA, while also recording information about single particle diameter. While ^siEVP^PRA uses molecular beacons for detecting RNA, SPIRFISH makes use of single-molecule fluorescence in-situ hybridization (smFISH) probes, which have improved signal to noise ratios and are more economical to be adapted as compared to custom molecular beacons. Importantly, SPIRFISH can be performed with commercially available technology and does not require in-house synthesis of custom surfaces for particle immobilization. Additionally, the protein and RNA labeling is a single-step process taking place concurrently at 37°C using 20% hybridization buffer, which facilitates smFISH probe binding without interfering with antibody binding. While the majority of the work done in our study utilizes tetraspanin capture antibodies, customized capture antibody chips are commercially available – allowing flexible capture options and analysis of a variety of particle types.

HIV virions served as an optimal particle to develop and validate the SPIRFISH technique, as infectious virions contain two copies of HIV genomic RNA and the HIV envelope protein gp120 at their surface; as such, we should be able to identify this population by RNA/protein colocalization (*27*, *29*). Additionally, they have been shown to be around 100-120 nm in diameter, which allows us to incorporate and investigate the size element for our analysis (*30*, *31*). Populations of HIV virions possess natural heterogeneity, due to inefficiencies in viral production that result in virus-like particles missing viral proteins or RNAs (*13*, *14*). These inefficiencies can be advantageous for the virus, as non-infectious virions can serve as decoys to soak up host defenses like neutralizing antibodies, amongst other functions (*57–59*). If infectious enveloped virions and host, non-viral EVs are thought of as bookends to a gradient of virus-like particles that exist between, deconvolution of these populations would prove useful (*13*). Finally, virologists have long desired a technique to examine properties of infectious virions without the influence of co-isolated non-viral or noninfectious particles.

Our initial SP-IRIS analysis of HIV demonstrated the usefulness of this single-particle technique even without the RNA component added by SPIRFISH; gp120 antibody labeling in combination with interferometric sizing revealed a novel population of larger, gp120+ particles present only in the HIV sample and primarily captured by a CD81 antibody. These are likely CD81+ HIV virions, although it cannot be concluded whether they are infectious virions from this analysis. Unfortunately, the gp120 antibody was imperfect and partially labeled the smaller population in the HIV sample and the non-viral EVs in the control sample. It is possible that the smaller particles being labeled in the HIV sample are smaller EVs carrying HIV-1 gp120 without other viral components; EVs carrying viral envelope proteins is a known phenomenon (*38*, *39*). The labeling seen in the EV control condition suggests, however, that it is at least to some extent non-specific binding of the gp120 antibody. Nevertheless, we believe that true gp120 staining of HIV virions can be differentiated from this non-specific background as it is 1.) coincident with a larger IM signature, 2.) brighter than the background by fluorescence intensity, and 3.) a population not present in the EV control. An introduction of IM and fluorescence intensity cutoffs within the analysis software could thus be used to reduce the contribution of background and focus in on the virion population for further colocalization analyses. Use of our SPIRFISH technique further deconvoluted the HIV sample with successful detection of single-particle HIV gRNA. Notably, the RNA fluorescence measured by SPIRFISH had low background and was shown to be specific and dependent on probe/target complementarity through use of non-specific control probes. SPIRFISH allows this single-particle RNA signal to be analyzed for colocalization with CD81, gp120, and particle size measured by interferometry (IM). This allows for discrimination between noninfectious virions (CD81+/gp120+/IM_high_/gRNA−) and infectious virions (CD81+/gp120+/IM_high_/gRNA+). Interestingly, only a small fraction of CD81+/gp120+/IM_high_ particles are gRNA+, supporting the literature that suggests a majority of released virions are noninfectious (*13*, *14*). Importantly, discrimination of infectious virions in this analysis uses only two of three available fluorescent channels afforded by the commercial SP-IRIS scanner.

To demonstrate the broad applicability of the SPIRFISH technique, we next attempted to detect specific tRNAs and their fragments in EVs. tRNAs have been found to be one of the most abundant extracellular RNAs; this includes tRNAs exported in EVs, where they often constitute a large proportion of the EV-internalized RNA molecules (*42*). Building upon this understanding, our investigation turned to the detection of tRNAs, particularly either full-length 5’ tRNA^Gly^GCC or it’s nicked forms and fragments. For tRNA detection, we focused on two cell lines: U2-OS, and Expi293F. The successful detection of these specific tRNAs within EVs derived from these cell lines stands as a testament to the robust capabilities of the SPIRFISH technique, showcasing its efficacy even when targeting very small RNA molecules. The tRNA smFISH signal was relatively low intensity and only detectable in a small fraction of captured EVs; however, these results should be contextualized. Weak signal could be explained by the low number of smFISH probes that can bind a single tRNA due to its short length. Likewise, low frequency could be explained by tRNA-containing EVs being rare. Alternatively, it is possible that the smFISH probes cannot bind to full length tRNAs due to tRNA secondary structure and lack of denaturing conditions during SPIRFISH, leaving only nicked tRNAs and tRNA fragments detectable. Given that full-length tRNAs are the most abundant species detected in U2-OS EVs (*41*), this could explain the low tRNA smFISH+ particle numbers. Despite low signal intensity and frequency, tRNA smFISH signal was still clearly detectable above background. This finding not only underscores the versatility of SPIRFISH in capturing diverse RNA targets but also highlights its potential utility in unraveling the intricate molecular landscape of extracellular vesicles.

Finally, we used SPIRFISH to detect mCherry mRNA loaded in engineered EVs. The engineered COURIER system seeks to provide two key functions: the export of barcodes to track cell populations, and the export of mRNA to allow one cell to “send” mRNA to another cell, where it can be expressed. Both applications rely on efficient and selective export of target RNA species. The successful specific detection of mCherry mRNA, its colocalization with structural EGFP, and the assessment of mCherry mRNA loading efficiency offer an example of how SPIRFISH can be used to determine the quality of engineered particles created for the therapeutic delivery of RNA. The SPIRFISH analysis presented here reveals a key quantitative parameter of the system: ∼14% of particles are loaded with cargo mRNA. This result provokes the question of whether different RNA cargo sequences or different exporter variants might produce different efficiencies of cargo loading. More generally, they suggest that it may be worth pursuing further engineering of the protein nanocages and RNA attachment sites to increase overall export efficiency.

Although our SPIRFISH technique is capable of co-detecting RNA and protein within a single particle to deconvolute particle populations, there are limitations that need to be considered. First, SPIRFISH analysis is currently limited to particles that are captured on the SP-IRIS microchip; which may or may not be representative of the bulk particle population depending on the capture antibodies used. We have found that the universal tetraspanin capture chips work well for most biological particles, although some particles can be tetraspanin negative. Microchips can be functionalized to capture the particles of interest, if prior information about surface molecules is obtained. Second, colocalization analyses are limited by the number of detection channels on the scanning instrument. Selected fluorophores on antibodies and smFISH probes must be compatible with the instrument and avoid spectral overlap, if possible. Third, detection of very low abundance particle-associated RNAs by SPIRFISH may be difficult as microchip capture spots can become oversaturated if high numbers of particles are added to compensate for low RNA abundance. Oversaturation results in overly bright signal from fluorescent antibodies targeting protein; additionally, the interferometric sizing becomes skewed and inaccurate on oversaturated chips even without fluorescence. Fourth, we do detect background fluorescent signal within the channel used for SPIRFISH that could be originating from the SP-IRIS microchips themselves as we have controls to check for specificity of smFISH probes. Lastly, the SPIRFISH technique could likely be further optimized. Currently, our protocol includes smFISH probe hybridization at 37°C for 1 hour, using 20% hybridization buffer, with 200 ug of probes per chip. Changes in the hybridization temperature, incubation time, type of buffer used, amount of probe added, and use of denaturing conditions for reducing target RNA secondary structure may improve the SPIRFISH signal magnitude and signal to noise ratio. We recommend that smFISH probe pools are individually validated before use in SPIRFSH. Additionally, most EV samples we have tested require a permeabilization step prior to probe hybridization to allow the smFISH probes to access the EV interior. On the contrary, our HIV virions had similar levels of gRNA smFISH signal regardless of whether we permeabilized or not (data not shown) that could imply that virions are naturally permeable for short smFISH probes.

We foresee our SPIRFISH technique having broad applications in the nanoparticle field. Virologists can use SPIRFISH in order to study the surface characteristics of infectious enveloped RNA viruses with unprecedented detail. Most previous analyses of virions and EVs were performed in bulk and not at single-particle resolution, thus not reflecting population heterogeneity (*49*, *50*). Single-virion analyses were performed using low-throughput techniques not capable of extrapolating conclusions to population level statistics, and seldom analyzed protein/RNA colocalization (*60*, *61*). With SPIRFISH, two fluorescent channels can be dedicated to detection of infectious virions (one for the viral entry protein, and one for the genomic RNA) while the third channel can be used to detect any surface or internal protein of interest. SPIRFISH thus allows focused analysis of a subpopulation (infectious virions in this hypothetical) without influence of the remaining population (noninfectious virions and EVs). In the field of EV therapeutics, EVs are engineered via producer-cell DNA transfection to package multiple proteins and RNAs (*6*, *62*). The presence of these RNA/protein cargos within the same EV is usually essential for the EV’s therapeutic effect. When multiple plasmids are used, some cells and the EVs they release will not possess the full suite of engineered molecules; SPIRFISH can aid in determining the efficiency of therapeutic EV production by calculating the percentage of single EVs that have all of the engineered elements via colocalization analysis. Theoretically, SPIRFISH could be used to detect two or even three independent RNA molecules within a single particle. Longer single-particle mRNAs could be assessed for integrity by designing two unique smFISH probe pools, each linked to a different fluorophore, that target opposite ends of the mRNA molecule. Codetection of both RNA signals by SPIRFISH would suggest an intact mRNA, while either signal independently would suggest an mRNA fragment was packaged. Further advancement of SP-IRIS technology would benefit the SPIRFISH method; this could include the addition of a fourth laser (Ultraviolet or near-infrared) to allow more markers to be detected during colocalization analysis.

## Materials and Methods

### Experimental Design

The objective of this study is to validate and put to use a new high-throughput single-particle analysis technique that can co-detect RNA and protein (SPIRFISH). To this end, we design smFISH probe pools against various RNA targets found within different nanoparticles. We then apply SPIRFISH analysis to various nanoparticles, including HIV virions, to demonstrate the technique’s utility. We hypothesized that SPIRFISH analysis would allow us to deconvolute nanoparticle subpopulations through RNA/protein co-detection.

### Cell culture

PM1-BaL cells, a suspension HuT-78 derivative T-cell lymphoma line (HIV Reagent Program #3038) chronically infected with the CCR5-tropic HIV-1 BaL strain (NIH GenBank Accession, #AY713409), were grown in T175 flasks (60 mL) at 37°C and 5% CO_2_. The cells were grown in RPMI-1640 supplemented with 10% FBS, 1% penicillin/streptomycin, 10 mM HEPES, and 2 mM L-glutamine. After supplementation, RPMI was sterilized by 0.22 μm vacuum filtration. Uninfected PM1 cells were grown under the same conditions. TZM-Bl HIV-1 reporter cells were grown in RPMI-1640 supplemented with 10% FBS and 1% penicillin/streptomycin. Cells were passaged biweekly. U2-OS cells (ATCC, #HTB-96) were grown in DMEM + 10% FBS in a T75 flask until 80% confluency, washed 3X, and then incubated in serum-free media (MEGM, Lonza) for EV collection. Bovine pituitary extract or antibiotics included in the media formulation were not added. Expi293F™ cells (Gibco, #A14527) were maintained in Expi293™ Expression Medium (Gibco, #A14351) in vented shaker flasks (Nalgene™ Single-Use PETG Erlenmeyer Flasks with Plain Bottom, #4115-1000) on a shaker platform maintained at 125 rpm in a humidified 37°C incubator with 8% CO2. Cell densities were maintained per manufacturer instruction, and used between passage 1 and 30 after the initial seeding.

### HIV-1 Virus Collection – Differential Ultracentrifugation (dUC)

Active viral production was confirmed by p24 ELISA testing of cell culture supernatant. Cell suspension was removed from the flasks during biweekly passaging and centrifuged in 50 mL conical tubes at 2000g for 5 minutes at 4°C to pellet cells. The virus-containing conditioned media (CM) was removed from the cell pellet and pooled before being frozen at −80°C to await further processing. Once 600 mL of CM had been obtained from 4 passages, the CM was thawed and centrifuged at 2000g for 20 minutes at 4°C to pellet remaining cells and large cell debris. The supernatant, clarified CM (CCM) was removed from the 2K pellet. The 2K pellet was collected by resuspension in 3 mL PBS. The CCM was then centrifuged at 10,000g for 20 minutes at 4°C to pellet apoptotic bodies and other large EVs. The supernatant (CCM+) was removed and the 10K pellet resuspended in 3 mL PBS. The virus-containing CCM+ was then passed through a 0.22 μm vacuum filter to remove residual apoptotic bodies. The resultant ultra-clarified CM (UCCM+) was then centrifuged at 100,000g for 90 minutes at 4°C using a Thermo Fisher AH-629 (36 mL polypropylene UC tubes) swing-bucket rotor (k factor 242) in a Sorvall wX+ Ultra series 80+ ultracentrifuge (max. acceleration and deceleration) in order to pellet the EV/virion mixture. The EV/virus pellet was resuspended in 3 mL PBS, and aliquoted into 30x 100 μL aliquots which were stored until use at −80°C. The 100K supernatant was also collected. The same workflow was used to collect extracellular vesicles (EVs) from uninfected PM1 cells as a control. To collect a media-only control, 60 mL of the supplemented RPMI media was processed using the above workflow, and the 100K pellet resuspended in 0.3 mL PBS.

### EV collection

U2-OS EVs: EVs were separated from conditioned media (CM) at the time point of 24 h, as previously described (*41*). The CM was centrifuged twice at 2,000g at 4°C and EVs were isolated by size-exclusion chromatography (SEC) using legacy qEV-original 70 nm columns (IZON), as per the manufacturer’s instructions, using 1x PBS as the mobile phase. EVs were concentrated by ultrafiltration (using 10 kDa MWCO filters) and stored at −20°C for no more than 20 days.

Expi293 EVs: Cells were removed from the CM by centrifugation at 1,500g for 5 minutes at 4°C. The supernatant was then centrifuged at 10,000g for 30 minutes. The supernatant was then centrifuged at 100,000g for 90 minutes at 4°C (max. acceleration and deceleration) using Sorvall wX ultra 80+ centrifuge with AH-629 (36 mL) swing bucket rotor (k factor 242). Pellet were re-suspended in 750 µL 1x DPBS, were aliquoted into 50 µL, and stored in Protein LoBind tubes (Eppendorf) at −80°C.

### p24 ELISA

p24 ELISAs were performed to confirm PM1-BaL viral production and to confirm enrichment of HIV-1 virions in the 100K pellet of the dUC procedure. ELISA was performed using the PerkinElmer Alliance HIV-1 p24 Antigen ELISA kit (#NEK050B001KT). Briefly 0.5 mL of unconcentrated CM or 1 μL of concentrated virus from the 100K pellet diluted with 899 μL PBS were mixed with 10X 5% triton-X following the manufacturer protocol and then serially diluted. Samples were then added in duplicate to the ELISA plate, and incubated at 4°C overnight. The following day, the plate was washed and measured on a BioRad iMark™ absorbance plate reader. Absorbance at 490 nm results were compared to a recombinant p24 standard curve.

### Western Blot

The HIV-1 BaL virus was prepared for immunoblotting with 2X Laemmli buffer under reducing conditions. The virus was loaded into a 4-15% TGX 18-well 30 μL stain-free gel at 100ng p24 per well. The Spectra multicolor broad range protein ladder from Thermo Fisher (26634) was also loaded to serve as a size reference. Proteins were separated by SDS-PAGE at 100V for 80 minutes. Proteins were then transferred to PVDF membranes using the Thermo Fisher iBlot2 system default P0 protocol. Membranes were blocked with 5% milk/0.1% Tween PBS and then probed overnight with antibodies against HIV-1 gp120 (Novus-Biologicals, #NBP3-06630AF488, 1:500), bovine serum albumin (Invitrogen, #MA5-15238, 1:1000), and HIV-1 p24 (Abcam, #Ab63913, 1:1000). The following day, membranes were washed and probed with secondary antibodies (Santa Cruz Biotechnology, #SC-2357 for rabbit primaries and #SC-516102 for mouse primaries, 1:5000). HRP signal was developed using Thermo Fisher SuperSignal™ West Pico PLUS Chemiluminescent Substrate (#34578) and imaged using a Thermo Fisher iBright FL1500 imaging system.

### Nano Flow Cytometry Analysis

The various samples obtained from the HIV-1 BaL production process (unprocessed material, 2K pellet, 10K pellet, 100K pellet, and 100K supernatant) were thawed and diluted 1:100 (unprocessed, 2K, 10K), 1:500 (100K pellet), and 1:10 (100K supernatant) in PBS. The samples were then loaded into and run on a Flow Nanoanalyzer from NanoFCM to obtain particle size and concentration data. The concentration calibration was performed using NanoFCM Quality Control Nanospheres and size calibration using 68-155 nm silica nanosphere mix (NanoFCM, #S16M-Exo). EVs from uninfected PM1 cells (collected from the 100K pellet) were run at 1:1000 dilution in PBS.

### Transmission Electron Microscopy

Media control, PM1 EVs, and HIV-1 BaL samples were diluted 1:10 into PBS and UV-inactivated for 20 minutes using a UV-crosslinker. Samples were then deposited onto 400 mesh carbon-coated TEM grids (EMS, ultra-thin) and negatively stained with Uranyl Acetate (1%, aq). TEM grids were imaged with a Hitachi 7600 TEM at 80kV with an AMT XR80 CCD.

### TZM-Bl HIV Reporter Assay

TZM-Bl cells were plated at 100,000 cells/200 µL/well in a 96-well flat bottom plate and allowed to adhere for 24 hours. The following day, the concentrated HIV-1 sample was thawed and serially diluted into RPMI from 500 ng p24 to 3.90625 ng. One group of the serially diluted HIV-1 doses was then UV-inactivated for 20 minutes using a UV-crosslinker, while the other group was left alone. RPMI was removed from the TZM-Bl cells and replaced with diluted virus-containing RPMI of varying p24 doses and +/− UV-inactivation. After 24 hours of virus incubation, the cells were lysed and transferred to white 96-well plates. Lysates were then assayed for viral infection induced firefly luciferase activity using the Promega Luciferase Assay System (# E1501) on a Thermo Fisher Luminoskan Ascent luminometer.

### Single-Particle Interferometric Reflectance Imaging Sensor (SP-IRIS) analysis of HIV-1

All SP-IRIS experiments used Tetraspanin capture chips from NanoView Biosciences (now called Leprechaun Exosome Human Tetraspanin Kit, from Unchained Labs, #251-1044) and an ExoView R100 instrument. Chips were pre-scanned prior to sample addition. Media control (1:50), PM1 EVs (1:100), and HIV-1 BaL (1:50) samples were diluted into ExoView incubation solution before 20-minute UV-inactivation. 50 µL of diluted sample was then added to the center of each pre-scanned capture chip and incubated overnight to allow particle capture. The following day, unbound particles were removed through various washes following manufacturer protocol, manually and/or using ExoView Chip washer. Chips were then incubated with fluorescent antibodies against CD63 (CD63-CF647, from ExoView kit), CD81 (CD81-CF555, from ExoView kit), and HIV-1 gp120 (gp120-AF488, from Novus Biologicals, #NBP3-06630AF488). Antibodies were diluted in a blocking buffer provided in the kit. Chips were then incubated for one hour in the dark on a microplate shaker at 430 rpm. After incubation, unbound antibodies were removed by additional washes. Finally, chips were dried and scanned interferometrically and fluorescently using the ExoView R100 instrument. Single particle count data were normalized across different chips by background subtraction; the average particle count for any defined parameter (fluorescence or interferometric) from the triplicate control mouse IgG capture spots was subtracted from the corresponding particle count on the tetraspanin capture spots (CD9, CD63, CD81).

### HIV-1 BaL genomic RNA and other target single-molecule fluorescence in-situ hybridization (smFISH) probe set design

The DNA sequence corresponding to the genomic RNA sequence of the HIV-1 strain BaL was obtained from the HIV sequence database (https://www.hiv.lanl.gov/content/index). A set of Oligonucleotide probes, each 20 nucleotides in length were designed to be complementary to the entire length of the genomic RNA sequence using Stellaris Probe Designer tool from Biosearch Technologies. The probes were synthesized with modified 3’ amino group and equimolar concentration of the probes were pooled for *en mass* conjugation with Texas Red (TR) [Invitrogen, #T6134] or Alexa Fluor-647 fluorophores, and then purified via high pressure liquid chromatography (HPLC) using previously published protocol (*63*). A set of oligos targeting a plant RNA (DCL4) and the Chikungunya virus (CHIKV) NSP2 RNA were also prepared in the similar manner. Additional probes were created to target the 5’-tRNA^Gly^GCC and mCherry. A list of all probe sequences is provided in the supplementary table (Data S1).

### Cellular validation of smFISH labeling of HIV-1 BaL genome

The hybridization protocol for suspension cells was followed to image the cells as described previously (*21*). Briefly, uninfected PM1 or chronically infected PM1-BaL cells were pelleted from suspension and washed with PBS. The cells were then fixed with 4% paraformaldehyde (4% PFA) by resuspension of the cell pellet. After 10 minutes in the fixation buffer, cells were pelleted again before resuspension in 70% ethanol. Cells were pelleted and subsequently washed with 20% wash buffer (containing 2X saline sodium citrate solution (Ambion, #AM9763), 20% formamide (Ambion, #AM9342/44) and 2 mM ribonucleoside-vanadyl complex (New England Biolabs, # S1402S)). The cell pellet was suspended in 100 μL of 20% hybridization buffer (containing 20% (vol/vol) formamide, 2 mM ribonucleoside–vanadyl complex, 10% (wt/vol) dextran sulfate (Sigma, #D8906), 1 μg/μL yeast tRNA (Invitrogen, #15401-029), 0.02% (wt/vol) ribonuclease-free bovine serum albumin (Ambion#AM2618), dissolved in 2X saline sodium citrate solution and containing 100 ng/μL of the smFISH probe set. The hybridization was performed in a moist chamber at 37°C overnight with gentle rocking. The cells were pelleted, and unbound probes were removed by serial washing of the pellet with 20% wash buffer four times. The cells were then pelleted and resuspended in wash buffer. The cells were suspended in antifade mounting medium containing DAPI and a drop of cell suspension was applied on a clean glass slide and a clean coverslip was placed on top of it making sure no air bubbles are formed. The edges of coverslip were sealed using clear nail polish. 100X oil objective was employed to capture images using a Nikon TiE Inverted epifluorescence microscope equipped with a Pixis 1024 b camera (Princeton Instruments) and Metamorph imaging software, version 7.8.13.0 (Molecular Devices). 16 Z-stack images, 0.2 mm apart were captured for each fluorescent wavelength using 1 second exposures. Images were processed using NIH developed ImageJ software.

### SPIRFISH - Combined SP-IRIS and smFISH analysis of HIV-1

ExoView chips were pre-scanned prior to sample addition. Media control (1:50), PM1 EVs (1:100), and HIV-1 BaL (1:50) samples were diluted into ExoView incubation buffer before 20-minute UV-inactivation. 50 µL of diluted sample was then added to the center of each pre-scanned capture chip and incubated overnight to allow particle capture. The following day, unbound particles were removed through various washes following the non-permeabilization manufacturer protocol. Chips were then incubated with fluorescent antibodies against CD81 (CD81-CF555, from ExoView kit) and HIV-1 gp120 (gp120-AF488, from Novus Biologicals, #NBP3-06630AF488), and 2 µL of smFISH probes (from 100 ng/µL smFISH probe stock). Antibodies and smFISH probes were diluted in 20% hybridization buffer. Chips were then incubated for one hour in the dark on a microplate shaker at 430 rpm, at 37°C. After incubation, unbound antibodies and unhybridized smFISH probes were removed by additional washes. Finally, chips were dried and scanned interferometrically and fluorescently using the ExoView R100 instrument.

### EV SPIRFISH with tRNA probes

ExoView chips were pre-scanned prior to sample addition. U2-OS EVs (1:5, or 1:2), and Expi293F EVs (1:2000) samples were diluted into ExoView incubation buffer and added to the center of each pre-scanned capture chip and left overnight at RT to allow particle capture. The following day, unbound particles were removed through various washes following the Chip washer cargo-protocol. Chips were incubated with fluorescent antibodies against CD81 (or Syntenin-1, from ExoView kit) and CD9 (from ExoView kit), and 2 µL of smFISH probes (from 100 ng/µL smFISH probe stock). Antibodies and smFISH probes were diluted in 20% hybridization buffer. Chips were then incubated for one hour in the dark on a microplate shaker at 430 rpm, at 37°C. After incubation, unbound antibodies and unhybridized smFISH probes were removed by additional washes. Finally, chips were dried and scanned interferometrically and fluorescently using the ExoView R100 instrument.

### EPN24-MCP EV Production and SPIRFISH with mCherry probes

HEK293T cells were plated on 10 cm dishes with 6,000,000 cells per dish, and co-transfected the following day with 10 mg of RNA exporter expression plasmid and 10 mg of cargo RNA expression plasmid using calcium phosphate. The RNA exporter EPN24-MCP-T2A-EGFP expression plasmid was pFH2.105 (Addgene, #205550). Cargo RNA expression plasmids for mCherry-MS2×8 and Cre-MS2×8 were pFH2.22 (Addgene, #205537) and pJAM1.52, respectively. Media was harvested 48 hours after transfection. Three such 10 cm dishes were prepared separately for each sample, then pooled. Pooled exporter particles were purified and concentrated approximately 500-fold by ultracentrifugation in a cushion of 20% (w/v) sucrose in PBS. Particles were stored at −80°CC until further use. For SPIRFISH, ExoView chips were pre-scanned prior to sample addition. EPN24-MCP Cre-mRNA (∼1:380) and mCherry mRNA (∼1:250) EVs were diluted into ExoView incubation buffer and added to the center of each pre-scanned capture chip and left overnight at room temperature to allow particle capture. The following day, unbound particles were removed through various washes following the Chip washer cargo-protocol (which includes fixation and permeabilization). Chips were then incubated with 2 µL of mCherry smFISH probes (from 100 ng/µL smFISH probe stock). smFISH probes were diluted in 20% hybridization buffer. Chips were then incubated for one hour in the dark on a microplate shaker at 430 rpm, at 37°C. After incubation, unhybridized smFISH probes were removed by an additional ∼30X washes (following the Chip washer cargo protocol). Finally, chips were dried and scanned interferometrically and fluorescently using the ExoView R100 instrument.

### Statistical Analysis

Data were exported from various instruments and graphed/analyzed using GraphPad Prism (version 10). Raw SP-IRIS and SPIRFISH data analyzed in the ExoView analyzer software were processed using default fluorescence cutoffs at 200 fluorescent units. The default interferometric size cutoff for the ExoView is 50 nm. Individual ExoView chip triplicate antibody capture lawns were visually assessed for QC issues (scratches or large artifacts) and in some cases excluded from the analysis. Each SP-IRIS and SPIRFISH experiment and condition was performed on 3 independent ExoView chips. For three-way comparisons performed in figures 2B and 4B, statistical significance testing was performed with ordinary one-way ANOVA with Tukey’s multiple comparisons test. For two-way comparisons in figures 6B and 6C, statistical significance testing was performed with unpaired two-tailed T-test. A p value of less than or equal to 0.05 was considered significant.

## Acknowledgements

We would like to thank the Johns Hopkins University School of Medicine Microscope Facility and Barbara Smith for assistance with electron microscopy of EVs and HIV virions.

## Funding

ION/ARPA Initiative: Payload Delivery, Ionis Pharmaceuticals (ZT, MB, KWW)

National Institutes of Health R01MH116508 (MBE)

the Allen Discovery Center, a Paul G. Allen Frontiers Group advised program of the Paul G. Allen Family Foundation UWSC10142 (MBE)

National Institute of Biomedical Imaging and Bioengineering of the National Institutes of Health R01EB030015 (MBE).

Chen Postdoctoral Fellowship Award (FH)

Burroughs Wellcome Fund Career Award at the Scientific Interface (FH)

National Science Foundation Award 2244127 (MB)

Universidad de la República CSIC 22620220100069UD (JPT)

## Author Contributions

Conceptualization: ZT, MB, KWW

Data curation: ZT, OG

Formal analysis: ZT

Funding acquisition: ZT, MB, KWW

Investigation: ZT, OG, ZL, AK

Methodology: ZT, OG, ZL, KWW

Resources: ZT, OG, AK, ZL, FH, MBE, JPT, MB, KWW

Visualization: ZT

Writing—original draft: ZT

Writing—review & editing: ZT, OG, AK, ZL, FH, MBE, JPT, MB, KWW

## Data and Materials Availability

All data are available in the main text or the supplementary materials, or upon request from authors. Materials are available upon request and with a completed material transfer agreement.

